# Online Reference Trajectory Adaptation: A Personalized Control Strategy for Lower Limb Exoskeletons

**DOI:** 10.1101/2021.06.21.449311

**Authors:** Mohammad Shushtari, Rezvan Nasiri, Arash Arami

**Affiliations:** Department of Mechanical and Mechatronics Engineering, University of Waterloo, Waterloo ON, Canada; Department of Mechanical Engineering, University of Alberta, Edmonton, AB, Canada; Toronto Rehabilitation Institute, University Health Network, Toronto, ON, Canada

**Keywords:** Exoskeleton Control, Rehabilitation, Trajectory Adaptation, Human-Exoskeleton Interaction

## Abstract

This paper presents a novel method for reference trajectory adaptation in lower limb rehabilitation exoskeletons during walking. Our adaptation rule is extracted from a cost function that penalizes both interaction force and trajectory modification. By adding trajectory modification term into the cost function, we restrict the boundaries of the reference trajectory adaptation according to the patient’s motor capacity. The performance of the proposed adaptation method is studied analytically in terms of convergence and optimality. We also developed a realistic dynamic walking simulator and utilized it in performance analysis of the presented method. The proposed trajectory adaptation technique guarantees convergence to a stable, reliable, and rhythmic reference trajectory with no prior knowledge about the human intended motion. Our simulations demonstrate the convergence of exoskeleton trajectories to those of simulated healthy subjects while the exoskeleton trajectories adapt less to the trajectories of patients with reduced motor capacity (less reliable trajectories). Furthermore, the gait stability and spatiotemporal parameters such as step time symmetry and minimum toe off clearance enhanced by the adaptation in all subjects. The presented mathematical analysis and simulation results show the applicability and effectiveness of the proposed method and its potential to be applied for trajectory adaptation in lower limb rehabilitation exoskeletons.

## I. Introduction

Each year, millions of people lose their mobility due to aging, stroke, and spinal cord injury (SCI). While the majority of stroke survivors can walk, many never reach a level of walking that allows them to perform daily activities [1]. Walking recovery is also known to be the top priority for individuals with incomplete spinal cord injury (iSCI) [2], i.e. the majority of SCI survivors. Therefore, several assistive devices and techniques have been developed to facilitate their recovery. Particularly, assistive exoskeletons made the rehabilitation process more accessible and efficient. For instance, affected individuals can benefit from high-intensity training sessions, which can boost their neuromotor control recovery [3]. These devices also have a huge potential in enabling individuals with motor deficits to perform more physical activities, preventing muscle atrophy [4], and improve their quality of life.

Several studies have recently presented novel control strategies for exoskeletons, ranging from force control [5] and trajectory control [6] to pure reflex-based controllers [7], [8]. It has been shown that a pure position control strategy is not effective for iSCI rehabilitation since it does not encourage users to contribute actively to the gait [9]. Accordingly, an effective control strategy should partially assist patients while allowing voluntary active movements and maximally engage their neuromuscular system [10], [11]. This strategy, also called Assist-as-Needed (AAN), promotes neuroplasticity and can lead to neural control recovery. The main challenge facing an AAN controller is to adapt itself to the user’s neuromuscular system capacity and intention. Such adaptation will optimize the user-exoskeleton interaction while involving the user in the movement control as much as possible.

Exoskeleton trajectory modification is one of the approaches which can tackle this challenge. For instance, in [12], the proposed controller adjusts the swing trajectory based on the joint positions at the late stance phase. Another framework is to adapt the trajectory to minimize the human-robot interaction force, exoskeleton’s applied torque, or tracking error [13–19]. For instance, [13] presented three different control strategies for minimizing the interaction force between the exoskeleton and the patient. Also in, [16], an AAN control strategy was proposed to adapt the swing phase trajectory by online minimization of applied force and position error.

Optimization-based trajectory adaptation methods rely on the user’s ability to generate cyclic and stable motions. Accordingly, any trajectory that minimizes the cost function is assumed to be an acceptable solution. However, iSCI individuals’ movements are not always cyclic or reliable; consequently, the AAN controller can lead to nonphysiological movement patterns [13].

In this paper, we propose a novel trajectory adaptation method, which instead of solely relying on the interaction force minimization, incorporates a secondary objective in the cost function that guarantees convergence to a cyclic and stable walking pattern. The proposed adaptation method accounts for subject-specific motor capacities. Our extensive simulations on the developed model of human walking show the effectiveness of this approach for a variety of lower limb impairment levels.

## II. Problem Statement

Human and exoskeleton dynamics in interaction with each other (Fig.1) is described as

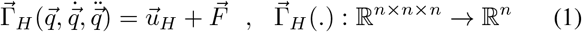

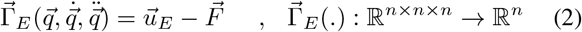

where 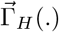 and 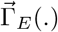 represent the passive dynamics of human and exoskeleton. We assume human and exoskeleton joints are coupled and 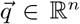 is the joint position of the exoskeleton and human (*n*: number of joints). 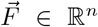 is the interaction force between human and exoskeleton. It is assumed that 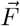 is directly measured using a force sensor. 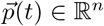 is the desired trajectory of human (human intention). 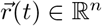 is the exoskeleton reference trajectory that will be adapted in interaction with human dynamics. 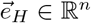 is the human error defined as 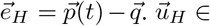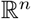 is the human applied torque which tries to minimize the human tracking error. Similar to human, exoskeleton tracking error and applied torque is considered as 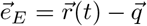 and 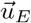, respectively. Exoskeleton reference trajectory 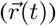 is modeled as the sum of initial trajectory 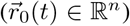 and a trajectory modification term 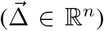. 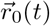 is a fixed *T* -periodic function with the period of gait cycle and 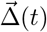 is left to be adapted. We parameterize 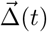 as a linear combination of basis functions as

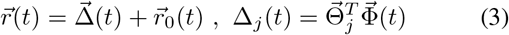

where *j* refers to *j*th joint. 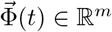 is a vector of *m* individual basis functions and 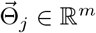 is the coefficient vector that is adapted to minimize the desired cost function. It is assumed that the basis functions are *T* -periodic, sufficiently smooth, and persistently exciting.

**Fig. 1.**
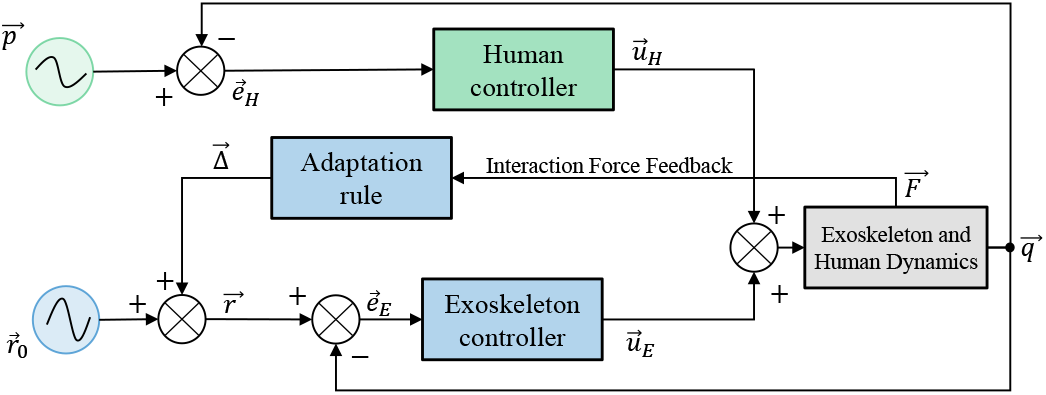
Human and exoskeleton dynamical interaction block diagram. The exoskeleton reference trajectory 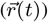 is adapted based on the feedback from interaction force 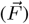. Human and exoskeleton controllers try to minimize their tracking error.

The proposed adaptation rule updates the bases coefficients at each joint 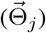 to minimize our suggested cost function (*J*) which is the sum of squared interaction force (*cost of interaction force*) and a weighted norm of the trajectory modification term (*cost of modification*), which penalizes deviation from the initial trajectory 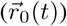.

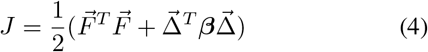

***β*** = diag(*β*_1_,*…*,*β_n_*) is a diagonal and positive definite matrix. For healthy users, we set ***β*** ≊ **0** which leads to interaction force minimization; i.e., the exoskeleton would be the follower of human motion. For users with lower motor capacity (e.g., those with iSCI) the user’s executed movement trajectories are not fully reliable to follow. Therefore, we consider *β_j_* > 0 to penalize the exploration of reference trajectory and facilitate maintaining a physiologically-plausible motion at those joints. The increment of *β_j_* depends on the motor capacity of impaired users at each joint; the higher the level of motor impairment, the larger *β_j_* should be.

Bipedal locomotion is generally an underactuated control problem and its stability is closely associated with the stability of zero dynamics [24]. Any conflict between the human and the exoskeleton controllers can act as a disturbance on the zero dynamics and adversely affect the gait stability and spatiotemporal parameters. In the next section, we propose a trajectory adaptation method that minimizes the conflict emerging in the form of human-exoskeleton interaction forces (Eq.4) and, therefore, enhances the gait quality. We also investigate the effect of our proposed method on i) gait stability, using the stability margin defined as the minimum distance between the zero moment point (ZMP) [25] and borders of the extended base of support [26], and on ii) spatiotemporal parameters of gait such as Step Time Asymmetry Index (STAI) [27], and minimum Toe Clearance (minTC) [28].

## III. Mathematical Analysis

### A. Adaptation rule extraction

To extract the adaptation rule for coefficient vector of the *j*th joint, we apply the gradient descent method to the cost function (Eq.4) as

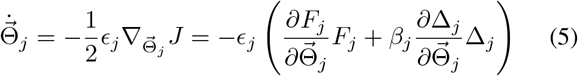

where *ϵ_j_* is the adaptation rate that tunes the speed of convergence. Without loss of generality, in the rest of this section, we drop the *j* index and follow our analysis at the joint level, therefore, 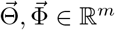, and the rest of the variables are treated as scalers.

Assuming a PD controller at each join of the exoskeleton yields 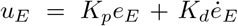 where *K_p_* and *K_d_* are the proportional and derivative gains, respectively. Therefore, from Eq.2, we obtain 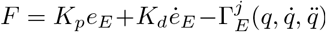 where 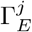 is the *j*th element of 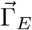 (we drop the *j* superscript from now on). According to Eq. 3, the exoskeleton error and its derivative are

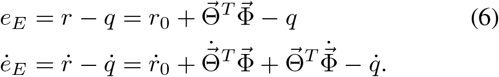

Therefore,

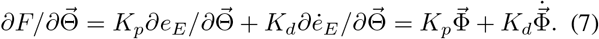

On the other hand, from Eq. 3, we have 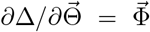. Substituting this expression and Eq.7 into Eq.5, we obtain the adaptation rule as

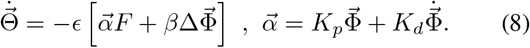

According to the Eq.8, only the user-exoskeleton interaction force (*F*) is needed to adapt the exoskeleton trajectory.

### B. Convergence analysis

To investigate the convergence and optimality of the adaptation rule, we substitute 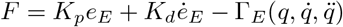, 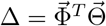, and Eq.6 into Eq.8, that yields

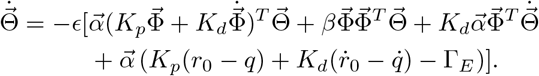

Performing some algebraic manipulations, the above equation can be rewritten as

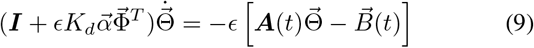

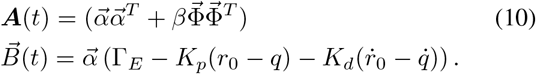

Choosing *ϵ* sufficiently small, we can ignore 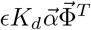 compared to the identity matrix and simplify the Eq.9 as:

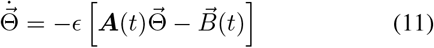

Since *r*_0_, 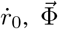, and 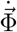 are periodic smooth and bounded functions of time, and *q*, 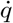, and 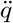 are bounded smooth and almost periodic [21] functions of time, we can compute the averaged adaptation dynamics using *General Averaging Theorem* presented in [22, pp. 413] as

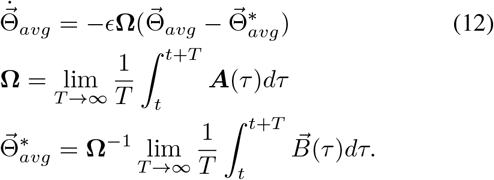

Since 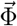 is persistently exciting, and 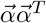 is positive semi definite, **Ω** is a positive definite matrix [23]. This guarantees the stability and convergence of the average adaptation dynamics (Eq. 12) toward 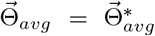. Therefore, the adaptation rule (Eq.8) is stable and convergent toward 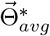 on average.

## IV. Simulator Development

To test the proposed trajectory adaptation method, we developed a dynamic gait simulator using exoskeleton and human musculoskeletal models in OpenSim. State feedback controllers were appropriately implemented within MAT-LAB (Fig.2).

**Fig. 2.**
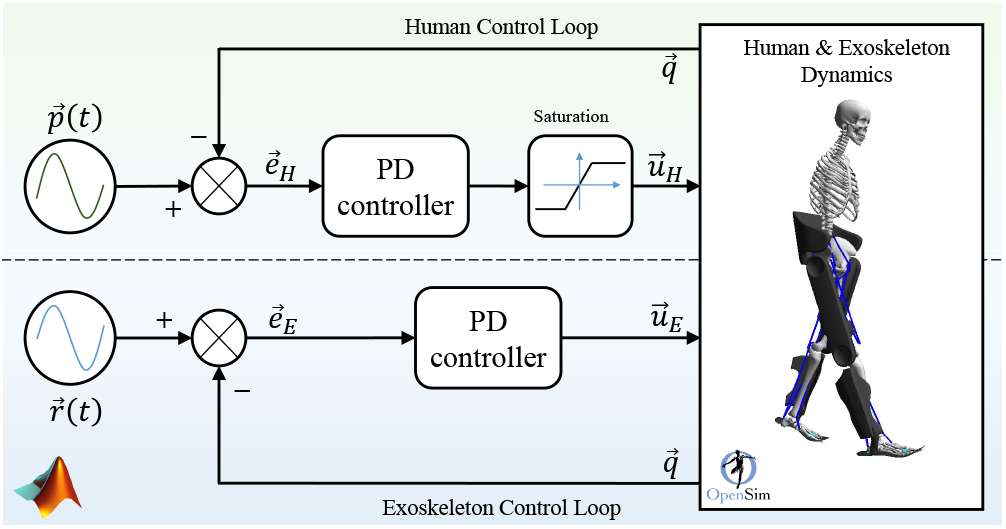
Dynamic walking simulator block diagram. The human and the exoskeleton controllers are implemented in MATLAB interfacing with the human and exoskeleton models which are defined in OpenSim environment along with ground reaction forces. The human tries to control his joint positions 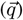 on his intended trajectory 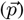 while the exoskeleton, with no information about the human intended trajectory, tries to minimize the tracking error between the exoskeleton position 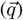 and reference trajectory 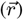. The saturation is added to the human control loop to simulate reduced motor capacity.

We utilized a 9-degree-of-freedom lower limb dynamical model in the sagittal plane presented in [29], with the ground reaction force (GRF) formulated using the model from [30], [31]. At each joint, the input torque is the sum of human 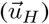 and exoskeleton 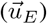 torque. Both human and exoskeleton controllers are assumed as PD controllers that apply torques based on the error from their own joint reference trajectories, which are 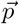 and 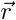, respectively.

### A. Simulator justification on healthy subject data

The human desired trajectories 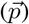 are fitted to the walking data (at 1.2 m/s) of a healthy adult (provided by OpenSim) with an eight-harmonic Fourier series. For the validation purpose, the exoskeleton model was excluded from simulation and its torques were considered to be zero 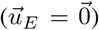. The human controller parameters are reported in Table I.

**TABLE I.** Exoskeleton and healthy human controller parameters.

| Parameter | Exoskeleton |  |  | Healthy Human |  |  |
| --- | --- | --- | --- | --- | --- | --- |
|  | Hip | Knee | Ankle | Hip | Knee | Ankle |
| $K_p$ [N/deg] | 34.9 | 22.68 | 34.9 | 22.68 | 13.96 | 31.41 |
| $K_d$ [Ns/deg] | 1.04 | 1.04 | 0.17 | 0.52 | 0.52 | 0.12 |
| Saturation Level [N.m] | — | — | — | 90 | 85 | 130 |

Fig.3 compares joint positions and torques of our simulator with those of real healthy human data. The real and simulated joint positions are almost equivalent with Pearson’s correlation coefficient (*R*) of 1. In addition, the torque profiles in all of the joints are comparable to the experimental torques in terms of sign and magnitude. Table II compares the *R* for each joint with two of the well-known biomechanical walking simulators ([33] and [34]). Our simulator exhibits a more realistic hip torque profile compared to the other simulators, while it has a comparable performance at the knee and ankle. Finally, the ground reaction force profiles of our model are similar to the results presented in [33] with *R* of 0.96 and 0.85 in *y* and *x* directions, respectively.

**TABLE II.** Simulated joint torques R-value compared to other works.

| Joint | Gayer | Van. Noot | Our work |
| --- | --- | --- | --- |
| $R_{Hip}$ | 0.45 | 0.53 | 0.76 |
| $R_{Knee}$ | 0.65 | 0.24 | 0.45 |
| $R_{Ankle}$ | 0.99 | 0.92 | 0.88 |

**Fig. 3.**
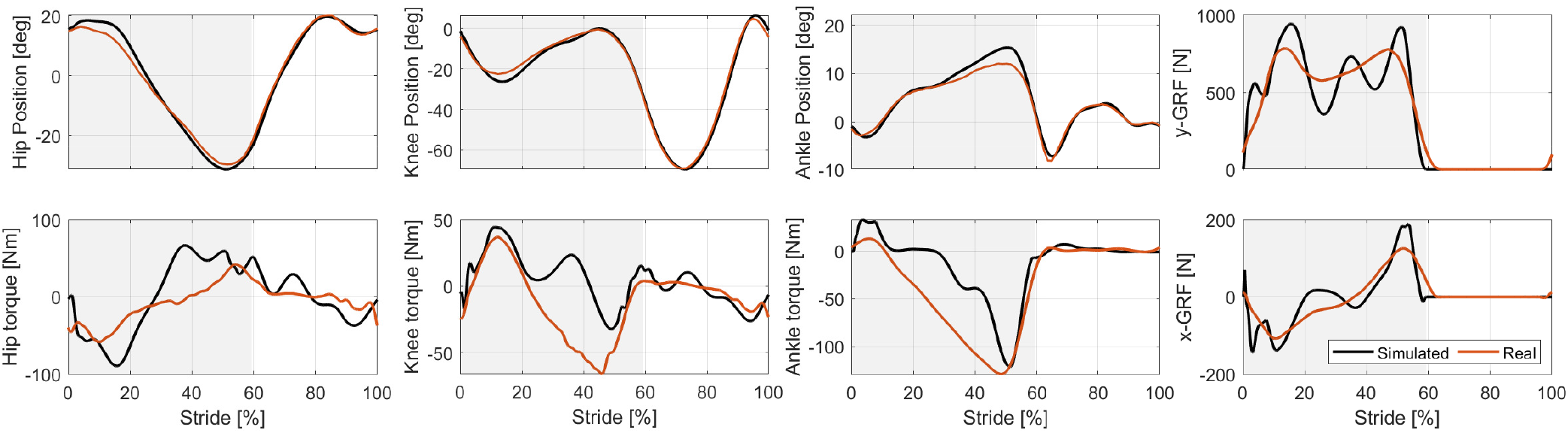
The comparison of the simulator and experimental data of a healthy subject (provided in [20]); the gray background indicates the stance phase. The position of the simulator joints closely follows the experimental human trajectory. The torque profiles’ sign and order of magnitude are also close to the ones of human torque profile such that the *R*-values for hip, knee, and ankle torques are 88%, 45%, and 76%, respectively. In addition, timing and order of magnitude for ground reaction forces are acceptable, and *R*-values for GRF in *y* and *x* directions are 96% and 85%, respectively.

### B. Simulation of assisted walking with an exoskeleton

Four different human subjects were simulated, one healthy and three with different motor impairments, subjects B, C, and D. To model different motor impairment levels, we considered a torque saturation block, imposing different saturation levels in the human control loop (Fig.2) and decreased the PD gains for the human controller with decreased torque range at more distal joints, which has resemblance to the reduced muscle recruitment in individuals with iSCI, who has ASIA score of B, C and D [32]. Parameters of mid-sever impairment was extracted from [32] indicating that iSCI with ASIA score C can apply between 24% to 36% of the knee extensor and 26% to 38% of the ankle plantar flexor torque that can be generated by healthy subjects. Accordingly, we limited joint torques of our mid-severely impaired subject at 30%, 35%, and 75% of healthy subject maximum joint torque at the ankle, knee, and hip joints, respectively. We assumed the two other impaired subjects to be less and more severely impaired (see Table III for simulated impaired subjects control parameters).

**TABLE III.** Relative Controller Parameters Of Impaired Subjects Compared To Healthy Subject (Table I).

| Subject | Saturation Level [%] |  |  | PD Controller Gains [%] |  |  |
| --- | --- | --- | --- | --- | --- | --- |
|  | Hip | Knee | Ankle | Hip | Knee | Ankle |
| **D** | 85 | 60 | 50 | 90 | 80 | 60 |
| **C** | 75 | 35 | 30 | 80 | 70 | 50 |
| **B** | 65 | 25 | 25 | 70 | 50 | 30 |

A three-harmonic Fourier series was fitted to the data of healthy human (provided in [20]) to define 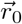. We perturbed the obtained Fourier coefficients so that the resultant exoskeleton initial reference trajectories considerably vary from the human intended trajectories 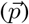, particularly at the hip and ankle. To maintain stability, the initial reference trajectory 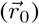 at the knee joint was chosen close to the human intended trajectory (Fig.4). Note that the exoskeleton reference trajectories cannot fully adapt to the human trajectories due to their fewer harmonics, more obvious at the ankle as more oscillations can emerge in comparison to other joints.

In the simulations, we assumed that the subject is walking on a treadmill. Therefore, there is no mismatch between the frequency of the human intended motion and the exoskeleton’s trajectories. In addition, for impaired cases, we have applied a soft constraint to regulate the orientation of the body w.r.t. the world frame; as motor-impaired individuals usually use walkers, walking sticks, and weight compensatory harnesses to be able to walk. The adaptation rule parameters are as follows: The basis functions 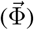 are the same as Fourier series bases, which are used for initial reference trajectory fitting. *ϵ* is also set to 6.5*e*-4, 3.2*e*-4, and 1.6*e*-4 for the hip, knee, and ankle trajectory adaptation, respectively, so that the basis coefficients 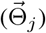 converge to their optimal values within 10 seconds. Adaptation was disabled in the first 10 seconds (8 steps). At *t* = 10*s* adaptation was enabled and simulations were continued for 60 seconds to ensure convergence and gait stability. We also tuned the ***β*** matrix according to the impairment of each subject and the saturation level at each joint Table IV.

**TABLE IV.** *β* values for simulation of each subject.

| Subject | $\beta$ | | |
| --- | --- | --- | --- |
|  | Hip | Knee | Ankle |
| **Healthy** | 0 | 0 | 0 |
| **D** | 3 | 91 | 304 |
| **C** | 6 | 152 | 760 |
| **B** | 15 | 304 | 1220 |

## V. Results and Discussion

Fig. 4 compares the adapted, initial, and intended trajectories during one gait cycle in four different simulated subjects. It indicates that, the exoskeleton reference trajectory at the hip joint closely follows the human intended trajectory after the course of adaptation in all subjects regardless of their impairment. The underlying reason is that the hip joint was the least affected joint in simulations. Thus, it can apply the sufficient torque required for the adaptation rule to decode the required trajectory modifications from interaction force. The knee and ankle joints, in contrast, were severely disabled in subject C and B. As a result, small deviation from the initial trajectories was observed. On the other hand, in the healthy subject and subject D, where those joints are less affected, we observe closer convergence to the human intention due to higher reliance on their performance.

**Fig. 4.**
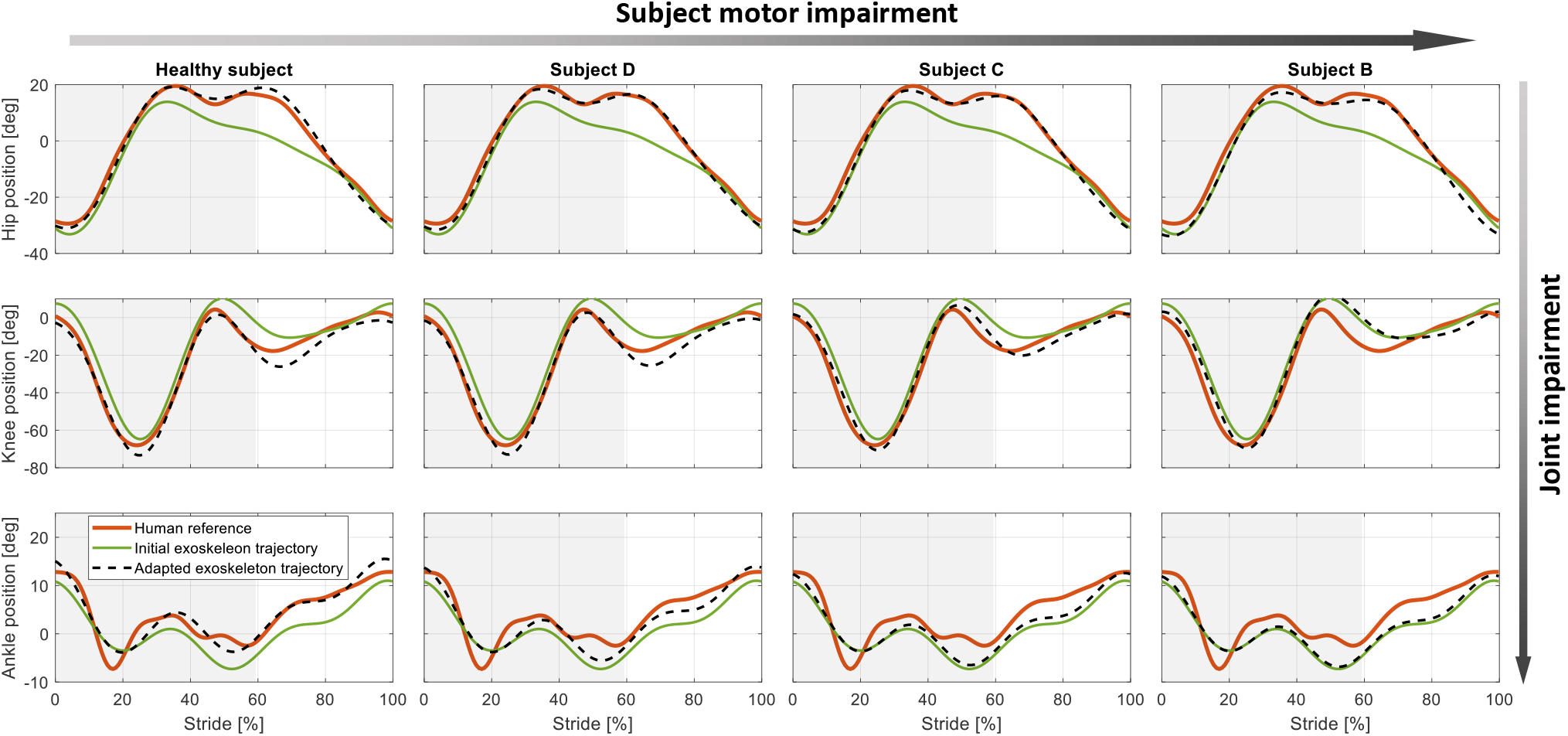
Exoskeleton reference trajectory adaptation at the hip, knee, and ankle joints (simulation results). The gray background indicates the stance phase. Each plot includes the human intended trajectory (in red), which is not known by the exoskeleton controller, initial exoskeleton trajectory (in green), and adapted exoskeleton trajectory (dashed lines). Subject B, C, and D are simulated with different reductions in motor capacity to generate the necessary joint torques for predefined intended motion according to Table III.

Fig. 5 illustrates the cost function, tracking error, and the trajectory modification during adaptation. The trajectory modification term and the interaction force costs are convergent while the former is increasing and the latter is decreasing in all cases. Those opposite competing trends provide an equilibrium point for the basis coefficients 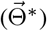, which is adjusted (personalized) by ***β*** matrix for each subject. Consequently, the interaction force cost decreased in the range of 43% for severely impaired subject (subject B) to 91% for healthy subject.

**Fig. 5.**
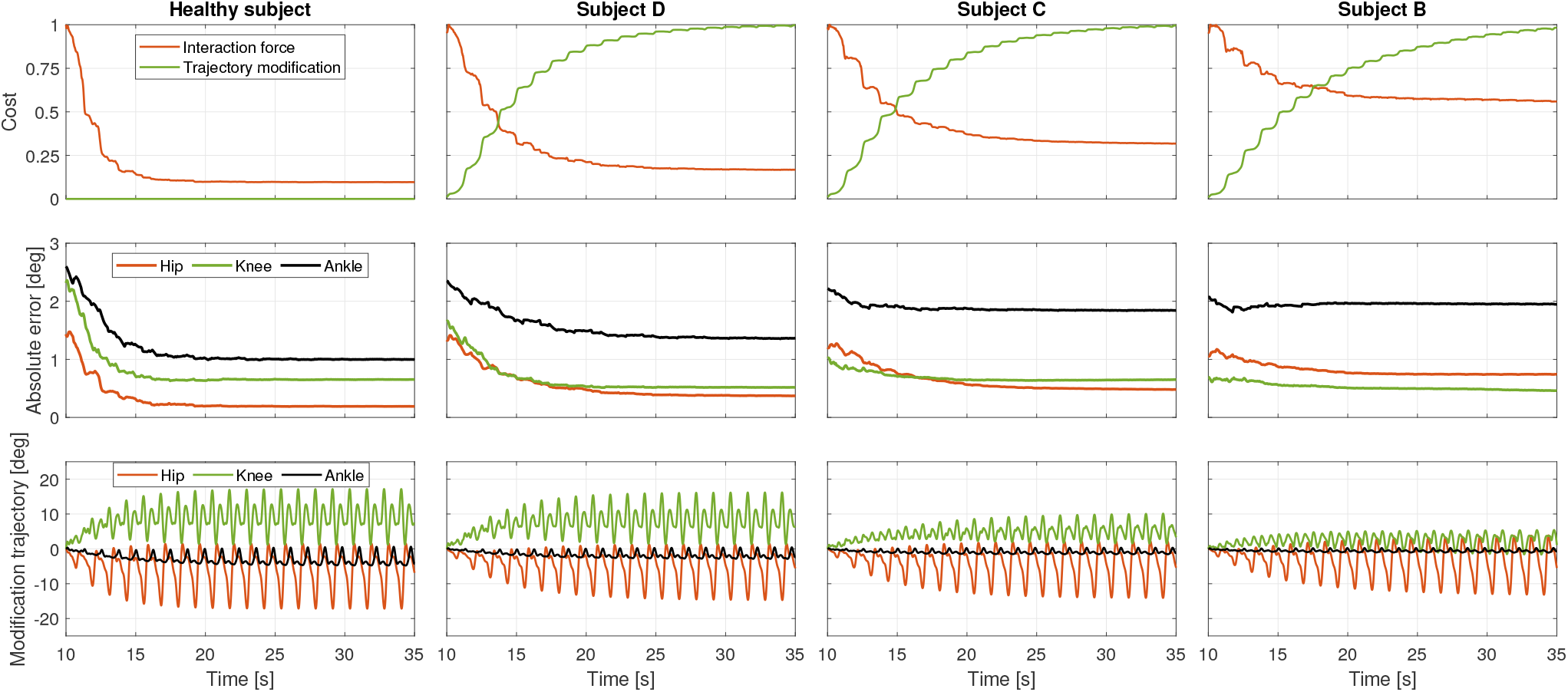
The adaptation performance (started at *t* = 10*s*) comparison in four different subjects. The first row presents the normalized moving average (the window length is *T* = 1.23*s*) of the trajectory modification and interaction force terms in cost function. The second row illustrates the moving average of exoskeleton absolute tracking error. Final row is the variations of the modification trajectories in course of adaptation. The interaction and the modification costs are convergent while the interaction cost is always decreasing. The tracking error is also decreasing in course of adaptation which shows the collaboration between trajectory adaptation and controller. Finally, modification trajectories are converged to a cyclic pattern, and by increasing the impairment of subjects, their range of variation is decreased.

Moreover, we observed that the healthy subject initially has a comparatively higher tracking error but eventually converges to a lower error since ***β*** = **0** does not penalize the trajectory modification. That is because the adaptation has higher flexibility to explore the parameter space, suggesting the active contribution of simulated user to the gait. As we move toward subject B, this observation becomes less pronounced due to the lower motor capacity of the subjects.

Adaptation of the exoskeleton reference trajectory leads to a drop in exoskeleton applied torque across all joints. The underlying reason is that the exoskeleton torque is the sum of dynamics compensation and interaction torques (based on Eq. 2) and while the former is almost fixed the latter is decreased. Besides, the decrease in tracking error indicates that adapted trajectories are more consistent with the dynamics of the human-exoskeleton system.

Fig. 6 shows the improved gait stability, step time symmetry, and an increased minimum toe off clearance, all as a result of trajectory adaptation which decreased the conflict between human and exoskeleton. Stability margin increased in the range of 60% for the severely impaired subject to more than 350% for the healthy subject. We noticed that the healthy subject has the lowest and the highest stability margin before and after the adaption, respectively, while the opposite trend was observed for the severely impaired subject. Adaptation has also eliminated the gait asymmetry which emerged more in the healthy subject. Finally, the minimum toe off clearance of the simulated subjects increased by about 40% on average.

**Fig. 6.**
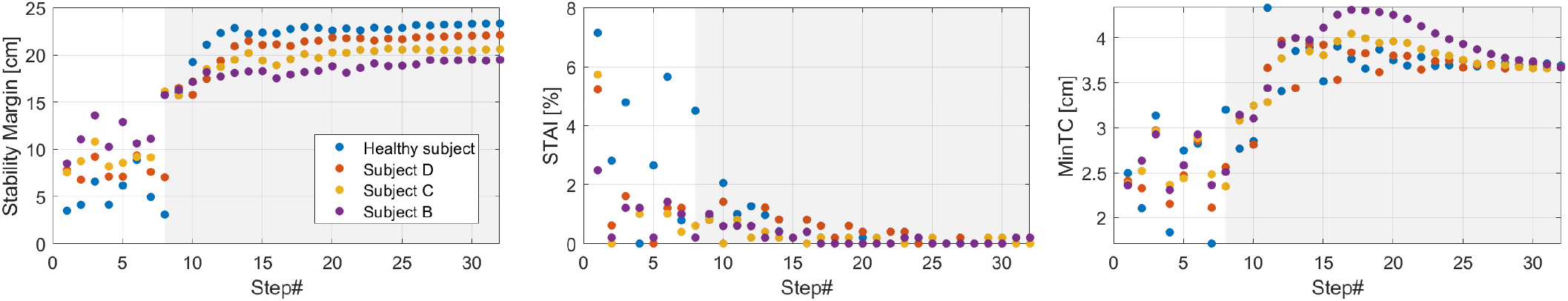
Adaptation effect on gait stability (left), step time asymmetry index (middle), and minimum toe off clearance (right) across subjects during the simulation. The gray area denotes the adaptation process started at *t* = 10*s* corresponding to step #8.

A limitation of the presented approach is the assumption that the frequency of human and exoskeleton reference trajectories are the same. While this can be held for treadmill walking, the assumption is not accurate in overground walking, requiring an adaptive approach to identify the gait frequency [35–37].

As mentioned in Section II, we did not include any adaptation capability for human desired trajectories. Therefore, a large mismatch between the initial trajectories of human and exoskeleton can lead o instability of the model. This issue is more pronounced in case of the simulated healthy subject as it is more capable of providing high torques to fight back the exoskeleton. However, in real world experiments, which is the next step of this work, human adaptation prevents severe mismatch and triggers the adaptation rule automatically to converge to new trajectory profiles such that both human and exoskeleton reference trajectories converge to their Nash equilibrium [38].

In this study, motor impairment was simulated by saturating and weakening joint controllers. We acknowledge that motor impairment can have other aspects which are not modeled here; however, our simulation environment provides a fair benchmark for testing the proposed trajectory adaptation. As a future work benefiting from the OpenSim engine integrated into Matlab, the human controllers will be reimplemented at the muscle level. This would enable us to simulate more complex impairments such as muscle flaccidity [39] or spasticity [40] in dynamical simulations.

## VI. Conclusion

In this paper, we presented an online trajectory adaptation technique based on interaction force measurement which has an adjustable capacity to converge to the human intended joint trajectories. We also developed a gait simulator that integrates MATLAB control and optimization toolboxes with the OpenSim musculoskeletal modeling frameworks. Using this toolbox, we applied the proposed adaptation method to a hypothetically healthy and three motor-impaired subjects, with similarities to iSCI patients with different levels of injury. The simulation results show that we can adapt the exoskeleton reference trajectories to the human intended trajectories in healthy subjects. On motor-impaired subjects, we also showed that the adaptation method is capable of restricting the trajectory adaptation in order to guarantee a stable, reliable, and rhythmic gait pattern. Exoskeleton trajectory adaptation also led to considerable improvement in gait stability and spatiotemporal parameters.

